# A functional chromogen gene *C* from wild rice is involved in a different anthocyanin biosynthesis pathway in *indica* and *japonica*

**DOI:** 10.1101/2020.08.24.264770

**Authors:** Weihua Qiao, Yanyan Wang, Rui Xu, Ziyi Yang, Yan Sun, Long Su, Lizhen Zhang, Junrui Wang, Jingfen Huang, Xiaoming Zheng, Shijia Liu, Yunlu Tian, Liangming Chen, Xi Liu, Jinhao Lan, Qingwen Yang

## Abstract

Accumulation of anthocyanin is a desirable trait to be selected in rice domestication, but the molecular mechanism of anthocyanin biosynthesis in rice remains largely unknown. In this study, a novel allele of chromogen gene *C*, *OrC1*, from *Oryza rufipongon* was cloned and identified as a determinant regulator of anthocyanin biosynthesis. Although *OrC1* functions in purple apiculus, leaf sheath and stigma in *indica* background, it only promotes purple apiculus in *japonica*. Transcriptome analysis revealed that *OrC1* regulates flavonoid biosynthesis pathway and activates a few bHLH and WD40 genes of ternary MYB-bHLH-WD40 complex in *indica*. Differentially expressed genes and metabolites were found in the *indica* and *japonica* backgrounds, indicating that *OrC1* activated the anthocyanin biosynthetic genes *OsCHI*, *OsF3H*, *OsANS*, *OsINS* and *OsANR* and produced six metabolites independently. Artificial selection and domestication of *C1* gene in rice occurred on the coding region in the two subspecies independently. Our results reveal the regulatory system and domestication of *C1*, provide new insights into MYB transcript factor involved in anthocyanin biosynthesis, and show the potential of engineering anthocyanin biosynthesis in rice.

**Author summary:** Accumulation of anthocyanin is a selection trait in rice domestication, whereas the mechanisms regulating the anthocyanin biosynthetic pathway in rice remain unresolved. Here, a novel allele of chromogen gene C from wild rice (*Oryza rufipongon*) was identified as a determinant regulator of anthocyanin biosynthesis. A key question is to what extent the involvement of the C1 gene can explain coloration variability of cultivated rice, where anthocyanin accumulation has been eliminated by artificial selection. Our results reveal the functional chromogen gene C from wild rice causes different coloration phenotypes, regulates various anthocyanin biosynthetic genes and produces different metabolites in *indica* and *japonica*. Artificial selection and domestication of the C1 gene in rice only occurs within the coding region of the two subspecies independently.

## Introduction

Most seed plants have different colors in different parts of seeds mostly due to the anthocyanin pigmentation [1]. Anthocyanins and pro-anthocyanins are a class of favonoid, as one of the largest groups of secondary metabolites, and are widely distributed in plants [2]. Anthocyanins contribute to not only multiple physiological roles in the responses of plant to biotic and abiotic stresses, but also commercial value of plant products. It was reported that various anthocyanin also have a profound impact on food quality beneficial to human health [3], which is of significant interest to both crop breeders and consumers.

Biosynthesis and accumulation of anthocyanin depend on the inherent genetic factors and external environmental factors. Inherent factors in anthocyanin biosynthesis include the structural and regulatory genes. Structural genes encode enzymes, include phenylalanine ammonia lyase (PAL), chalcone synthase (CHS), chalcone isomerase (CHI), flavonoid-3-hydroxylase (F3H), flavonoid-3’-hydroxylase (F3’H), dihydroflavanol reductase (DFR), and anthocyanidin synthase (ANS). The expression of these structural genes is controlled by transcription factors (TFs) and has been well characterized in a range of plant species [4–6]. A ternary MBW complex, comprising R2R3-MYB TFs, basic-helix-loop-helix (bHLH) TFs and WDR (WD-repeat) proteins, was believed to tightly regulate the common pathway of anthocyanins and pro-anthocyanins biosyntheses [7–10]. The regulating network of anthocyanin biosynthesis is well documented in maize and Arabidopsis but much less in rice. Sun et al. (2018) reported that a *C-S-A* gene system regulated anthocyanin pigmentation in rice hull. In this system, *C1* encodes a R2R3-MYB transcription factor and acts as a color-producing gene, and *S1* encodes a bHLH protein that functions in a tissue-specific manner. *C1* interacts with *S1* and activates the expression of *A1*, which encodes a dihydroflavonol reductase [11].

The R2R3-MYB TFs were often identified as determinants of variation in anthocyanin pigmentation and have been identified in many higher plant species [12–17]. In a previous study, *OsC1*, *OsRb* and *OsDFR* were identified as the determinants of anthocyanin biosynthesis in rice leaves [18]. The R2R3-MYB gene *OsC1* was previously isolated from cultivated rice through comparative mapping between rice and maize or according to the nucleotide sequence homology with known maize orthologues [19, 20], and was also cloned from cultivated rice using various methods recently [21, 22]. Several studies determined that *OsC1* acted as a chromogen gene and mainly functioned in apiculus and leaf sheath [23–27]. The apiculus, as the remnant of awn, maintains its color in some modern varieties and seems not to have undergone artificial selection during domestication [20]. The purple sheath trait was reported to be a morphological marker in rice [24], which can be easily observed at the seedling stage, and have often been used in the screening of the authentic hybrids. However, molecular function of *OsC1* was not fully understand, the exact genetic determinants for purple apiculus and leaf sheath in rice remain to be unraveled.

Rice ancestor wild rice (*Oryza rufipogon*) has accumulated anthocyanin in various tissues, most of *O. rufipogon* appear to have purple or red pigment mainly on awn, apiculus, stigma, pericarp, pulvinus, leaf blade, leaf sheath, internode and palea, but most cultivated rice have lost these pigments due to artificial selection. It had been demonstrated that *OsC1* was also a domestication related gene for the loss of pigments in cultivated rice [28]. Identification of the *C1* allele in wild rice is of great significance for understanding the origin and evolution of rice through investigation of the genes controlling color formation. In this study, a R2R3 MYB transcription factor was fine mapped in the wild rice using a set of chromosome segment substitution lines. The *C* gene from wild rice, *OrC1*, was cloned for functional analysis, transcriptom and metabolome profiling and was shown to be a functional allele for anthocyanin biosynthsis. Interestingly, we found that six anthocyanin metabolites were simultaneously regulated by *OrC1* in different genetic backgrounds. Our findings emphasize the importance and value of using wild relatives to uncover useful genes that have been lost during crop domestication in order to expand the genetic repertoire for rice domestication study and modern crop breeding.

## Results

### Fine mapping of *OrC1* using chromosome segment substitution lines

In our previous study, a set of Chinese common wild rice chromosome segment substitution lines (CSSL) was developed [29]. The donor parent, *O. rufipogon*, has a significant purple coloration in apiculus, leaf sheath, and stigma and black hull. The recipient parent, an *indica* variety 9311, has no purple pigment in the whole plant. In total 150 CSSLs was investigated under five environments (S1 Table) for purple coloration. One hundred twenty-one SSR and 62 InDel markers, which were polymorphic between the two parents and evenly distributed on all 12 chromosomes, were selected for genotyping (S2 Table). In the CSSL populations, the purple apiculus, leaf sheath and stigma were completely correlated, while the purple or black hull was independently segregated. Using SSR/InDel genotyping, one QTL related to purple apiculus, leaf sheath and stigma (hereafter named purple coloration trait) was identified under five environments and located near RM314 on chromosome 6. (Fig 1a)

**Fig 1.**
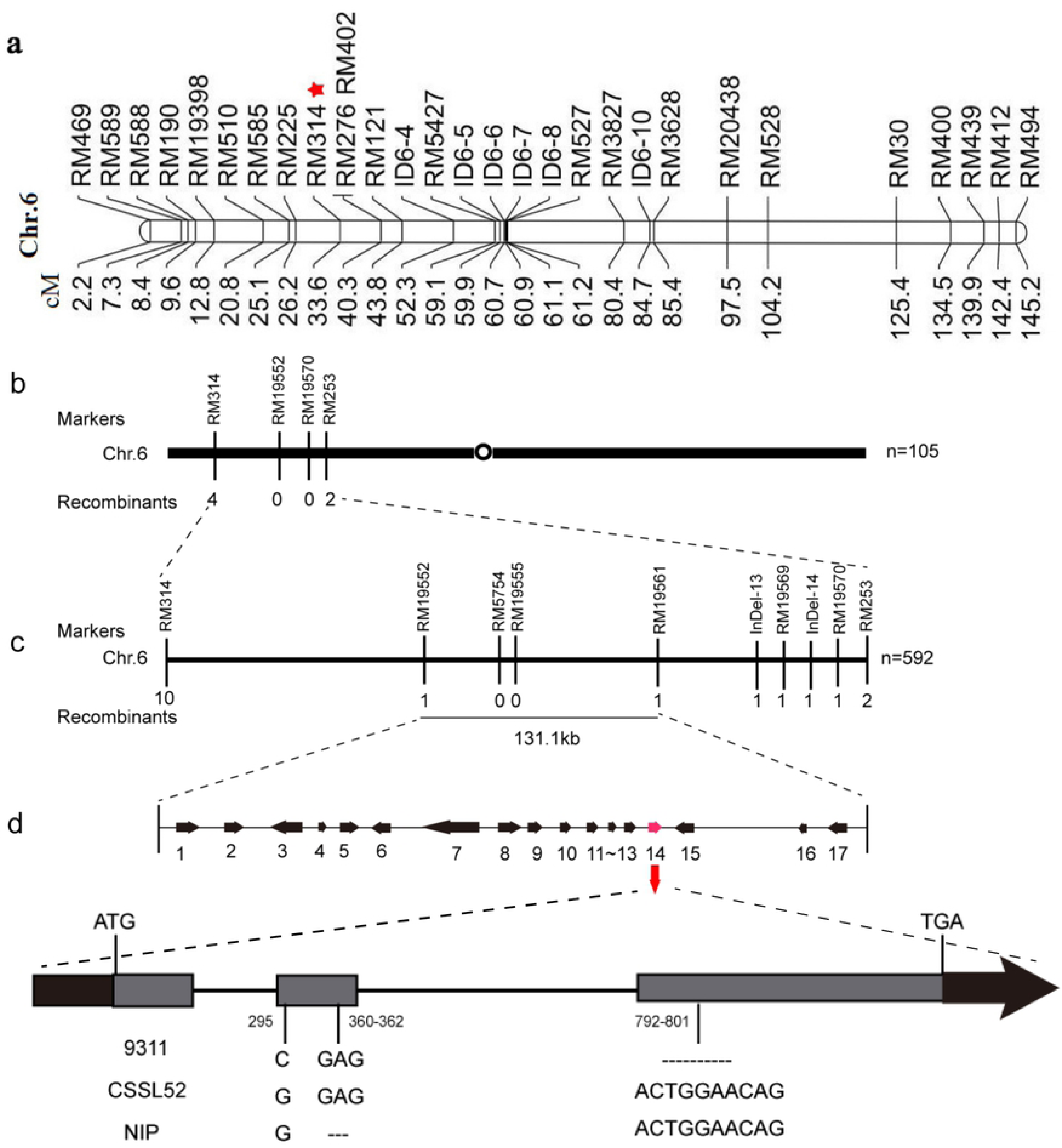
Fine mapping of *OrC1*. (a) Genetic map of chromosome 6 of CSSL52. The red star represents the marker closely linked with the QTL for the purple coloration trait. (b,c) The location of *OrC1* was narrowed down to a 131-kb interval between markers RM19552 and RM19561. The number of recombinants obtained is indicated under the marker names, and the number of individuals (n) used in mapping are shown on the left. (d) Seventeen predicted open reading frames (ORF) were identified in the fine mapped region. ORF14 encodes a homolog of *C1* gene. The deletions in 9311 and Nipponbare alleles were shown in (d).

One chromosome segment substitution line, CSSL52, which harbors this QTL and purple coloration trait, was selected for further fine mapping of purple genes. CSSL52 carried only one wild rice introgression segment and exhibited purple apiculus, leaf sheath and stigma (Fig 1a and Fig 2). We observed no significant difference in other agronomic traits such as plant height and heading date, between 9311 and CSSL52, indicating that the single introgression segment from wild rice was only involved in purple coloration. CSSL52 was back-crossed with 9311 to produce F_1_ plants, which showed purple apiculus, leaf sheath and stigma (Fig 2). The color of apiculus, leaf sheath and stigma in F_2_ population was completely correlated. A segregation ratio 3:1 of purple coloration to achromatic was observed in 1,964 F_2_ individuals (S3 Table), indicating that the purple coloration trait was controlled by a single dominant gene. Using F_2_ segregating population, this gene was narrowed down to a 131-kb region between markers RM19552 and RM19561 with recessive class analysis (Fig 1c), in which there were 17 predicted open reading frames (ORF) (S4 Table). This region contains an expressed gene (*LOC_Os06g10350*) encoding a R2R3 MYB transcription factor that has been reported as the rice homolog of maize *C1* gene [20]. Then we compared the nucleotide sequences of CSSL52 and 9311 and *japonica* Nipponbare across the fine-mapped region. Sequence analysis of *Os06g10350* revealed that 9311 contained a 10-bp deletion at the start of the third exon, and Nipponbare contains a 3-bp deletion in the second exon (Fig 1d). Therefore, we focused on *Os06g10350* as a candidate for purple coloration and named it as *OrC1*. Furthermore, a near isogenic line (NIL) of *OrC1* was developed using the positive F_2_ individual of CSSL52/9311.

**Fig 2.**
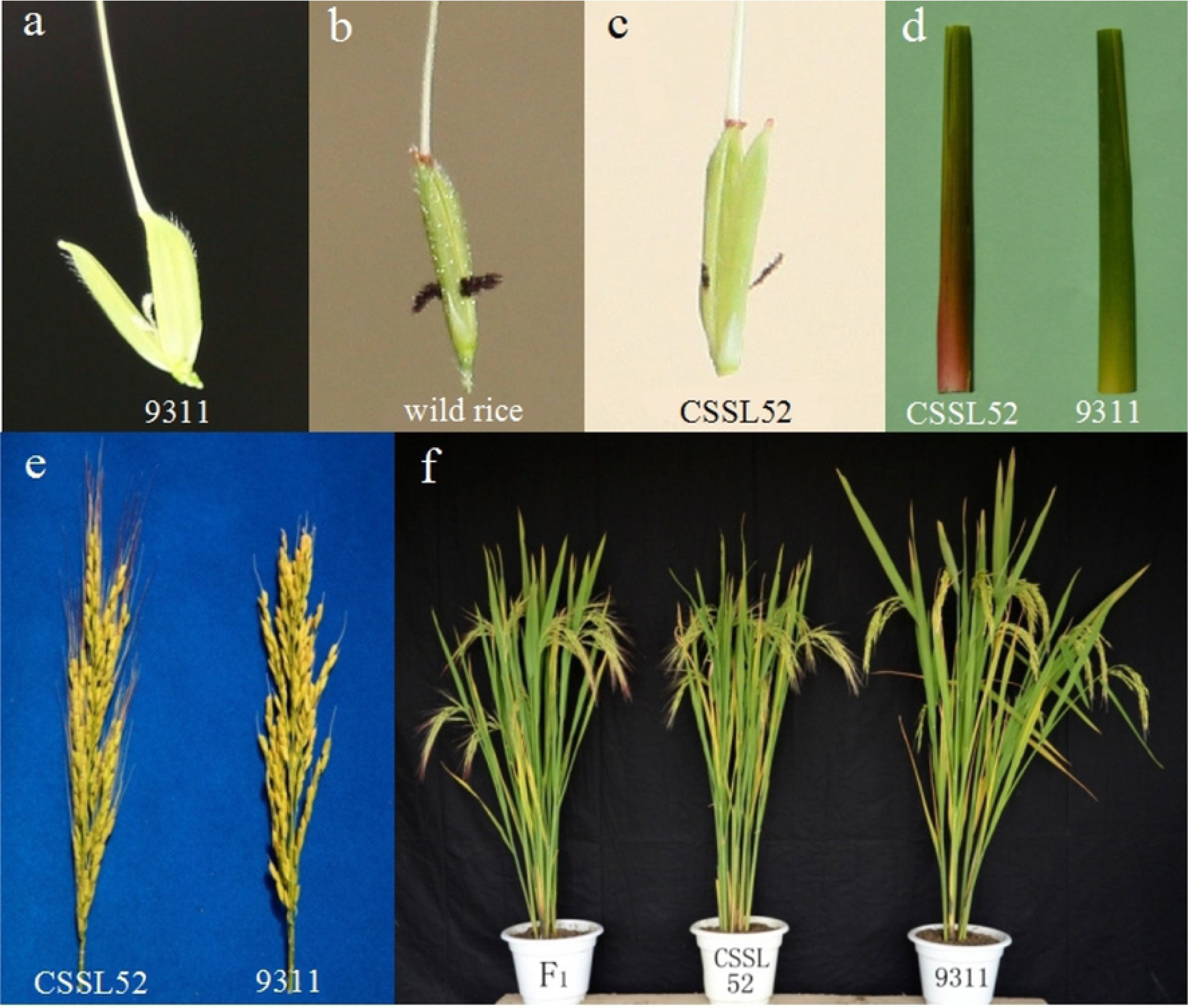
Phenotype of CSSL52, 9311 and F_1_ individual. (a-e) CSSL52 has purple apiculus (c), leaf sheath (d) and stigma (a-c) compared with 9311. (f) The F_1_ of CSSL52/9311 showed the same purple coloration trait as CSSL52.

### Characterization of OrC1 from wild rice

*OrC1* encodes a 272 amino-acid protein containing two highly conserved SANT domains, similar to the R2 and R3 motifs of R2R3-MYB transcriptional factors in plants. A phylogenetic tree was constructed using the amino acid sequences of *OrC1* and other anthocyanin biosynthesis related genes from several plant species. Phylogenetic analysis showed that *OrC1* exhibited highest homology to *OsC1* in cultivated rice and R2R3 MYB transcription factors in maize (Fig 3a). To investigate the expression patterns of *C1* in wild and cultivated rice, the levels of *C1* mRNA were quantified in different tissues including leaves, leaf sheaths, stems, and panicles of NIL-*OrC1* and 9311, at the booting stage and 3 days after flowering. No significant difference was detected between NIL and 9311. *C1* transcripts were detected at a high level in leaf sheaths, the lowest level of *OrC1* transcripts was detected in stems. Three days after flowering, the expression levels of *C1* in panicles was significantly increased in both NIL and 9311 (Fig 3b).

**Fig 3.**
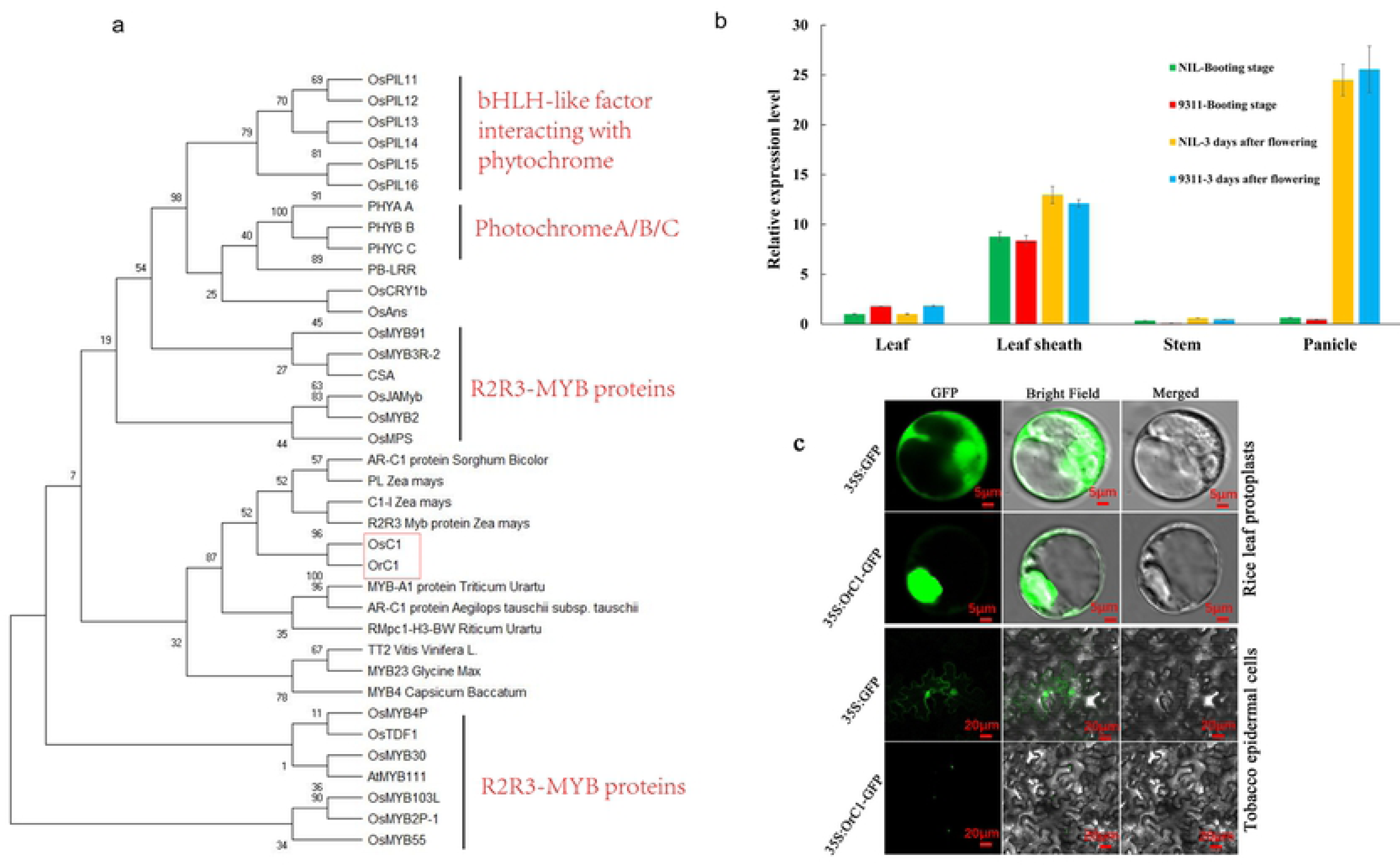
Characterization of *OrC1*. (a) Phylogenetic analysis of wild rice OrC1 and other R2R3-MYB proteins. (b) Expression pattern of *OrC1* gene in different tissues a two time points. (c) Subcellular localization of OrC1 in rice leaf protoplasts and tobacco leave.

To examine the subcellular localization of OrC1 protein, an OrC1:GFP fusion gene was generated and transformed into rice protoplasts and tobacco leaves under the control of the cauliflower mosaic virus 35S promoter. As shown in Fig 3c, the control GFP protein was distributed throughout the entire cell, whereas the OrC1-GFP fluorescent signals were exclusively localized in nucleus, consistent with the function of OrC1 as a transcription factor.

### Transcriptome analysis revealed differentially expressed genes between NIL and 9311

To further investigate the molecular mechanism of OrC1-mediated anthocyanin synthesis pathway, we performed a transcriptome analysis with the leaf sheath of 9311 and NIL at the seedling stage, because the purple color phenotype were most obvious at this stage. Three biological replicates were tested. The principal component analysis of the samples based on the number of fragments per kilobase of exon per million fragments mapped (FPKM) values showed that one replicate of NIL (OrC1-3) did not cluster with the other two (S1 Fig), so only two replicates were used for further analysis. In total, 2,388 differentially expressed genes (DEGs) were identified with the stringent criteria (|log_2_(FC)|>1, and FDR<0.05) (S5 Table). Of these DEGs, 1,225 were up-regulated and 1,163 were down-regulated in NIL compared with 9311. Furthermore, all DEGs were assigned to 98 KEGG. KEGG pathway enrichment analysis revealed that *OrC1* mostly affected the expression of genes involved in phenylpropanoid biosynthesis, diterpenoid biosynthesis, plant hormone signal transduction, and flavonoid biosynthesis (Fig 4a). To facilitate global analysis of gene expression, we investigated the possible mechanisms of significant DEGs by Gene Ontology (GO) analysis. The 2,388 DEGs were categorized into three main GO categories of biological process, cellular component, and molecular function. They were significantly assigned to biological process GO term, including cellular and metabolic processes. In terms of the cellular component GO term, they were mainly associated with cell, cell part, membrane and organelle. In respect to the biological process GO term, they were significantly involved in binding and catalytic activities (S2 Fig). The most enriched GO terms were metabolic process and catalytic activity.

**Fig 4.**
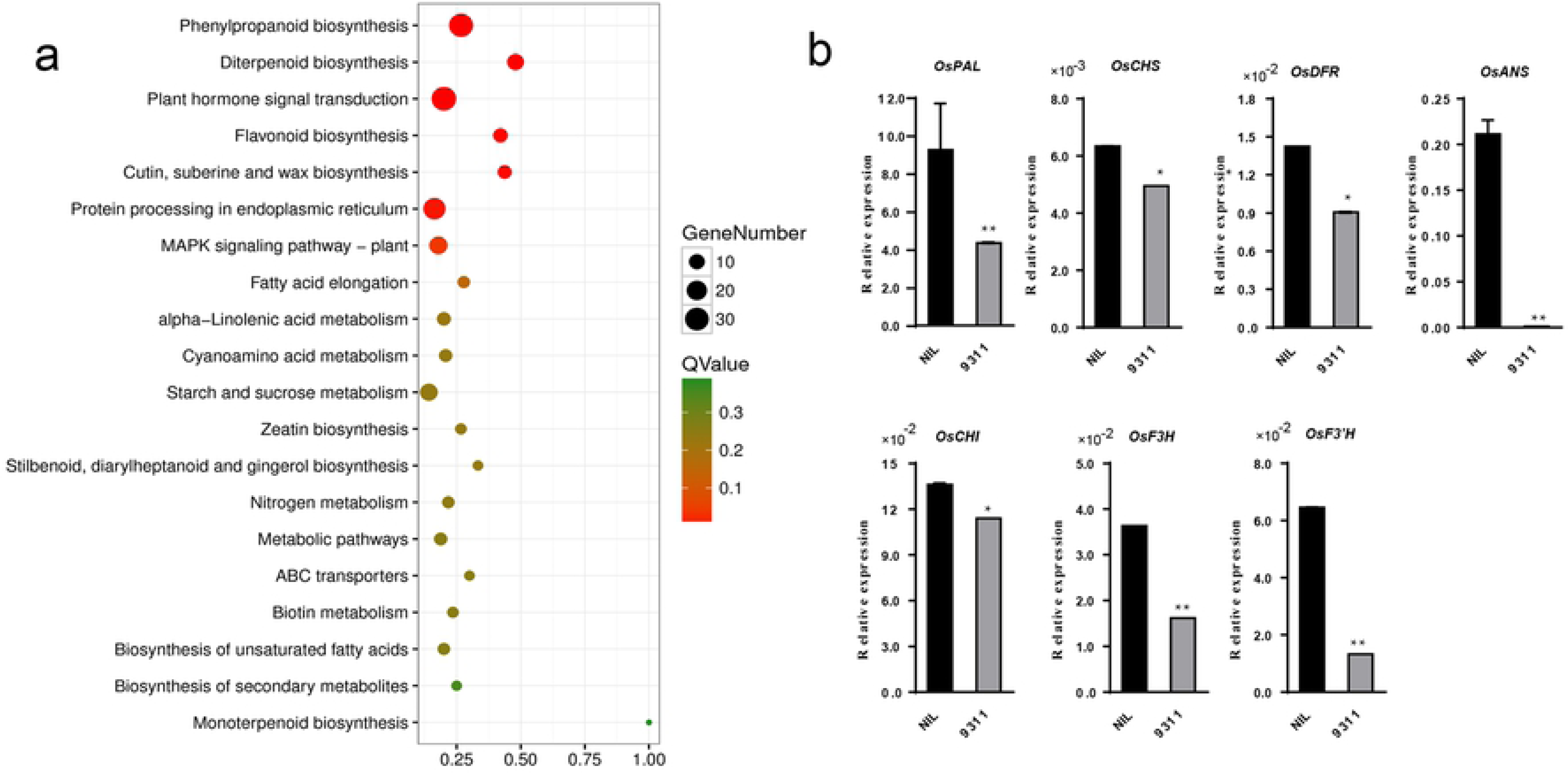
*OrC1* regulates gene expressions. (a) Top 20 significant enrichment pathway; (b) Relative expression levels of seven structure genes involved in anthocyanin synthesis determined by qRT-PCR. The values represent the mean ± s.d. *P < 0.05 and **P < 0.01 indicate significant differences in two-tailed Student’s t-tests. Three biological replicates. The actin gene was used for normalization.

Seven structural genes involved in the anthocyanin metabolic pathway were used for the validation of the sequencing results. The relative gene expression levels and FPKM (RNA-Seq) of the seven genes showed similar regulatory patterns in the NIL compared with 9311. A linear regression coefficient (R^2^) of 0.9711 was obtained between the results of qRT-PCR and RNA-Seq (S3 Fig). These consistencies validated the results of RNA-Seq analysis. As shown in Fig 4c. All seven anthocyanin biosynthetic genes were consistently up-regulated in the NIL compared with those in 9311, among them, expression levels of *OsPAL*, *OsDFR*, *OsANS*, *OsF3’H*, and *OsF3H*, were significantly increased in the NIL (Fig 4b).

Anthocyanin biosynthesis are largely influenced by some key pathways and MBW complex. According to transcriptome results, 116, 84, and 19 DEGs were detected in phenylpropanoid biosynthesis, MAPK signaling and flavonoid biosynthesis pathways, respectively (Fig 5a). At the same time, five *R2R3-MYB* transcription factors, six *bHLH* proteins and four *OsWD40* proteins were also found in DEGs (Fig 5b), including the cloned gene *Rc* (*Os07g0211500*) which was involved in proanthocyanin synthesis.

**Fig 5.**
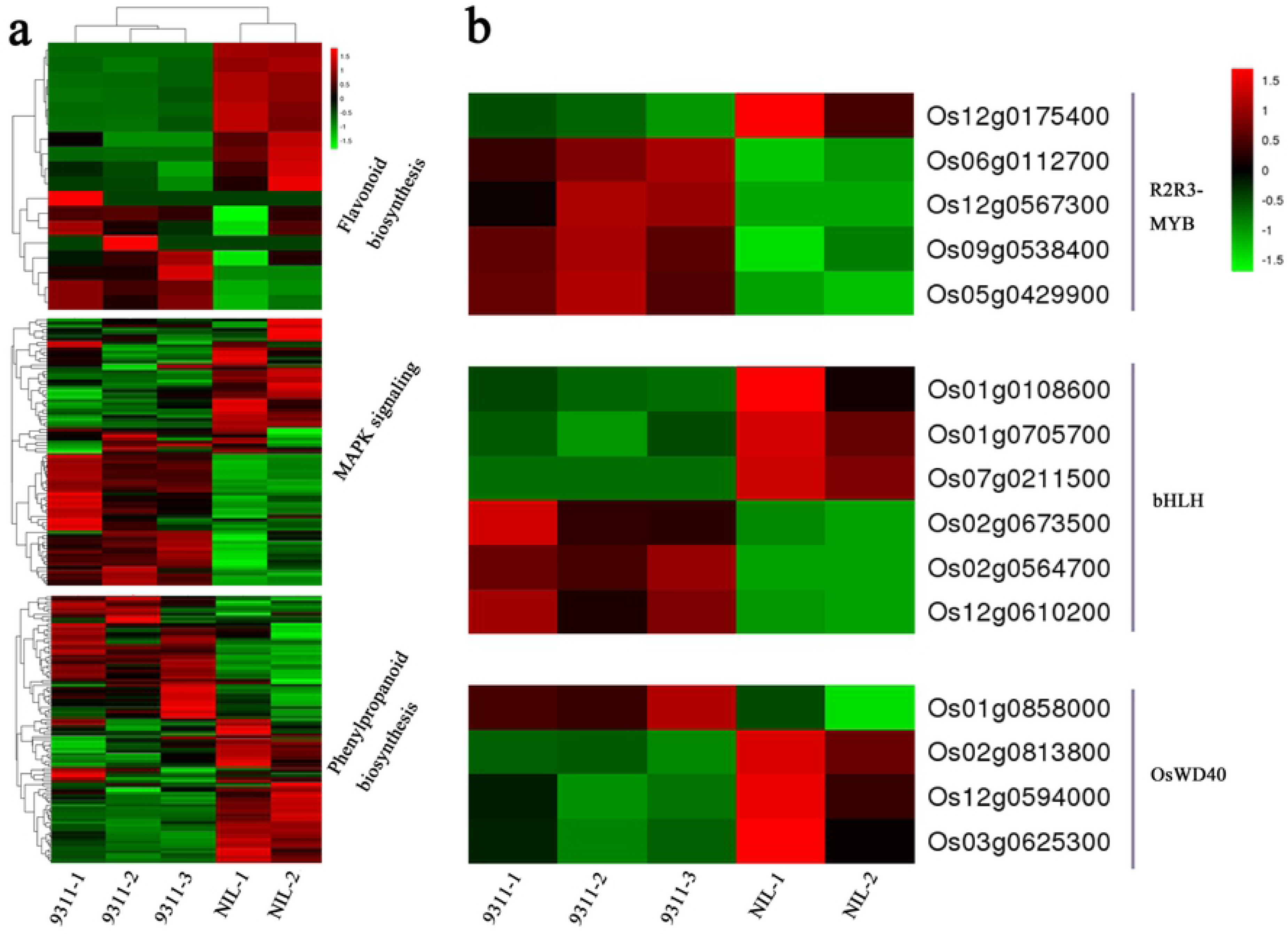
Heat map diagram of expression levels for anthocyanin biosynthesis related genes in (a) three KEGG pathways and (b) MBW complex. The heat map was drawn according to FPKM values. Columns and rows in the heat map represent samples and genes, respectively. The KEGG pathway or gene ID were displayed on the left. Color scale indicates fold changes in gene expression.

### Overexpression of OrC1 results in purple apiculus in transgenic Nipponbare

To explore the biological function of *OrC1*, its ORF was transformed into *japonica* cultivar Nipponbare with the 35S promoter. More than 20 independently transgenic lines carrying 35S:OrC1 (OE) were generated. The transgenic individuals were confirmed by PCR analysis with gene-specific primers. Ten positive transgenic lines were analyzed by qRT-PCR to determine the transgene expression level and two independent lines, OE1 and OE2, which differed in transcription levels were selected for additional analysis (S4 Fig). Under natural conditions, only the apiculus of T_1_ plants showed a purple pigmentation phenotype, the sheath was as the same as wild type Nipponbare. Purple apiculus of transgenic plants could be observed when panicles were exposed to sunlight at the initial heading stage. It turned into brown color at the wax ripeness stage. T_2_ transgenic plants showed a perfect segregation ratio 3:1 of purple or brown apiculus to normal. Both purple and brown apiculus could be easily distinguished at the fully ripen stage. There is no difference between the two *OrC1* overexpression lines, OE1 and OE2, and no other phenotypes were different between OE and Nipponbare except apiculus color (Fig 6a). Seven above mentioned structural genes were also detected by qRT-PCR, only *OsANS, OsCHI* and *OsF3H* showed higher expression level in OE lines than Nipponbare (Fig 6b). Other TFs, especially MBW complex, might involve in different pigmentation of Nipponbare and 9311, we compared other bHLH and WDR genes in NIL, 9311 and Nipponbare. All the gene sequences in NIL is exactly the same as 9311, which indicated that only one coloration gene *OrC1* in NIL was from wild rice. Comparison between 9311 and Nipponbare showed that six bHLH genes and three WDR genes have non-synonymous SNP (S5 Fig). Therefore, we predicted that these genes might have different functions in anthocyanin biosynthesis in different tissues.

**Fig 6.**
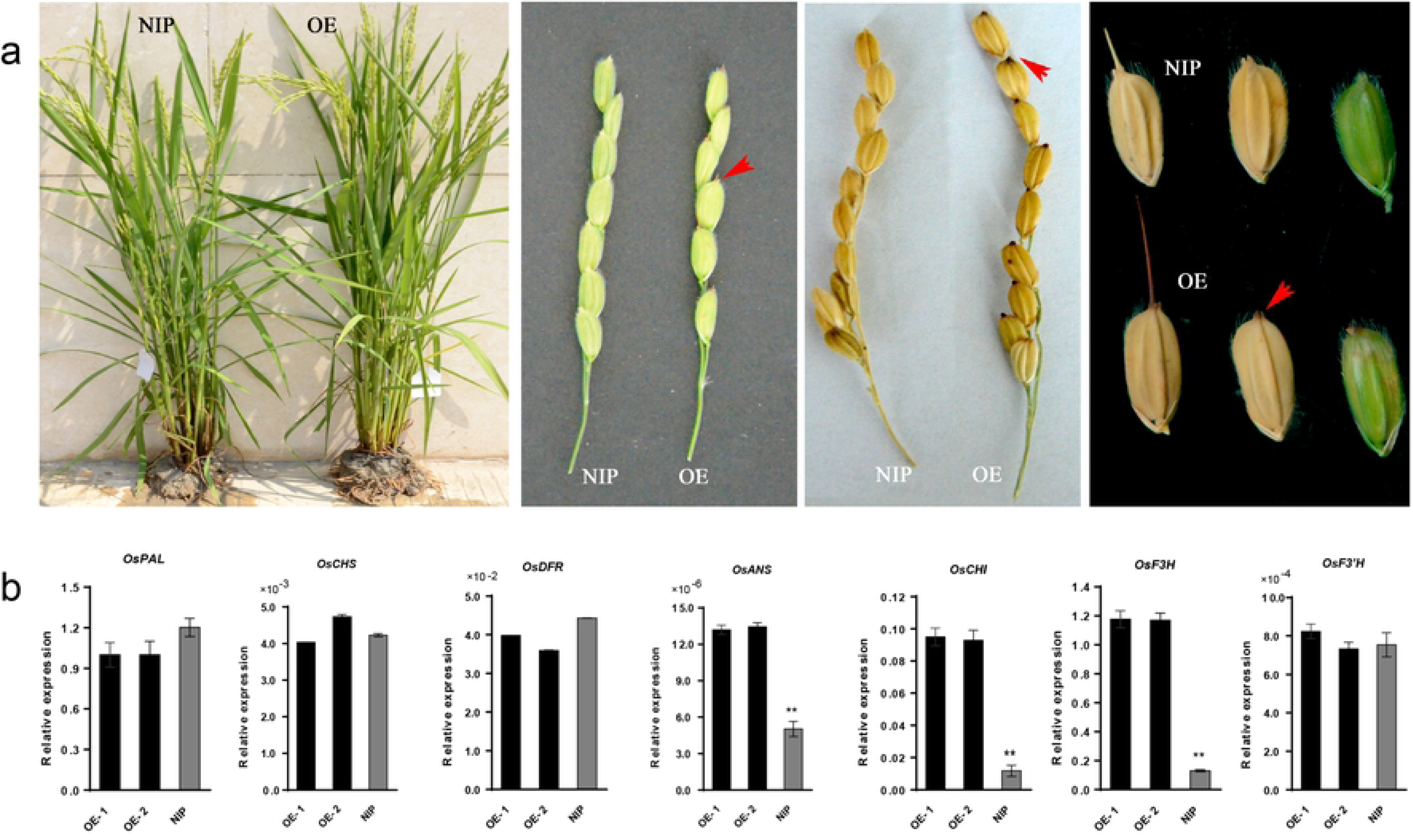
Overexpression of *OrC1* results in purple apiculus of Nipponbare. (a) Phenotypes of transgenic *OrC1* overexpression line (OE) and wild type Nipponbare (NIP) plants. The purple coloration was only found in the apiculus of transgenic plant. From left to right: whole plant, young spikelet, mature spikelet, mature and young seeds of NIP and OE. (b) The relative expression levels of seven structure genes determined by qRT-PCR in NIP and two OE. Three biological replicates. The values represent the mean ± s.d. *P < 0.05 and **P < 0.01 indicate significant differences in two-tailed Student’s t-tests. The actin gene was used for normalization.

### Identification of differential metabolites in the anthocyanin biosynthesis pathways

Liquid chromatography-electrospray ionization-tandem mass spectrometry analysis was used to quantify the anthocyanin content in leaf sheath of both NIL and 9311. A total of 28 metabolites of anthocyanins and pro-anthocyanidins were detected, and the relative contents for each metabolite were normalized before being subjected to downstream data analyses. Among them, 11 metabolites in NIL were significantly higher than that in 9311, two metabolites were higher in 9311 than that in NIL. For Nipponbare and transgenic plants, given the OE lines only have purple apiculus and no color in other organs, the leaf sheath and upper one third of the hull of two OE lines and Nipponbare were used for metabolite analysis. A total of 13 metabolites of anthocyanins were detected, and among them six metabolites in apiculus (hull) of OE were significantly higher than that in Nipponbare (Fig 7). In sheath, there is no significant difference between OE and Nipponbare. No difference was found in apiculus or sheath between the two OE lines.

**Fig 7.**
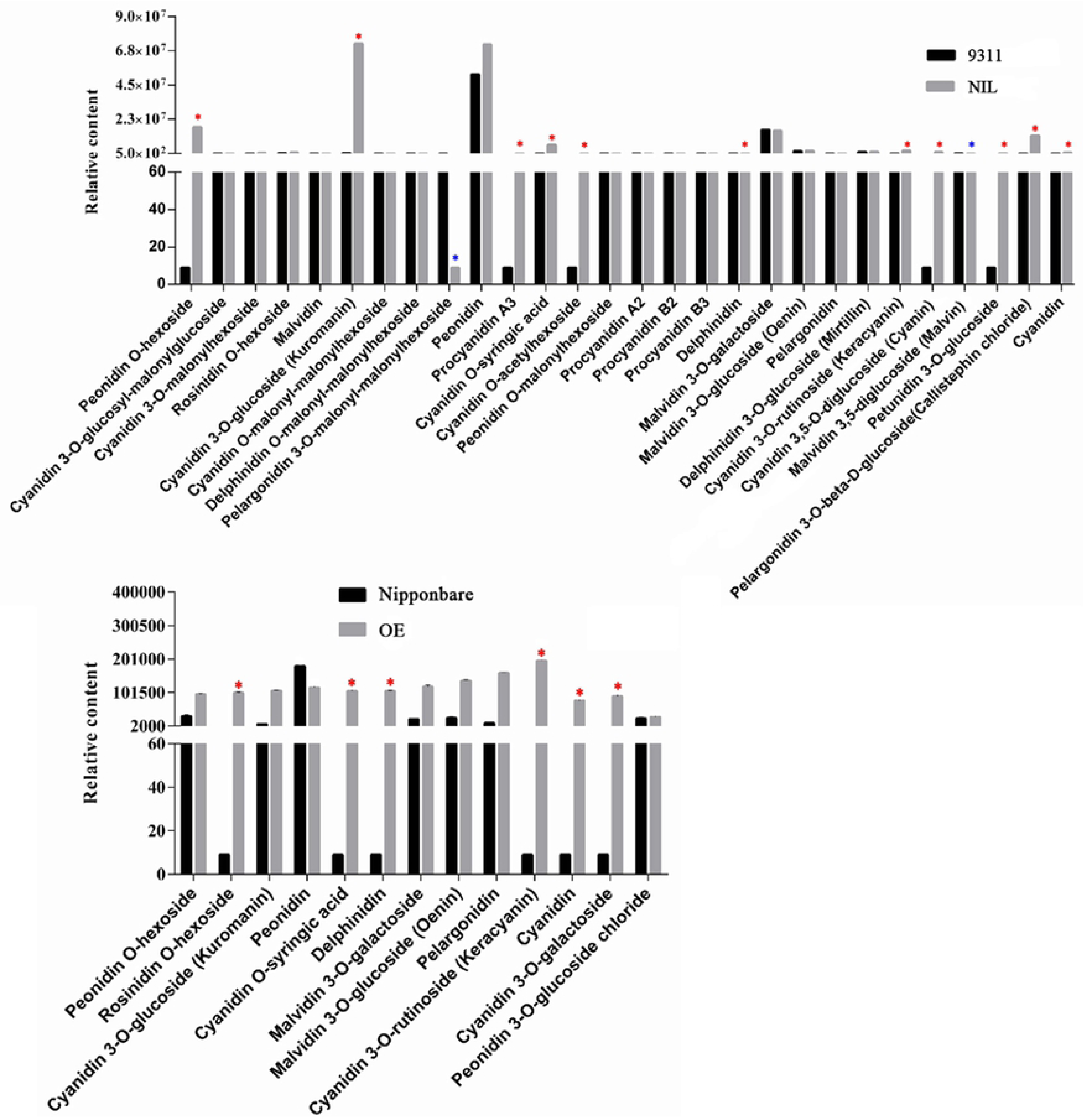
Identification of the differentially accumulated metabolites between NIL and 9311 (above), and between OE and Nipponbare (below). Red * indicates a significant increase at fold change ≥2, and blue * indicates a significant decrease at fold change ≤0.5.

The differentially accumulated metabolites (DAM) between pairs of samples (NIL-Sheath vs 9311-Sheath, OE-Hull vs Nip-Hull, OE-Hull vs OE-Sheath, OE-Sheath vs Nip-Sheath) were determined based on the variable importance in projection (VIP)≥1, and fold change ≥2 or ≤0.5 [30–31]. Comparative analysis of the four groups of DAMs revealed that six common anthocyanin metabolites were differentially accumulated in NIL-Sheath vs 9311-Sheath, OE-Hull vs Nip-Hull, and OE-Hull vs OE-Sheath, including rosinidin, delphinidin and four cyanidins (S6 Table). And all of them are up regulated in NIL compared with 9311, and in OE lines compared with Nipponbare.

Three genes were involved in delphinidin and cyanidin biosynthesis pathway in the flavonoid biosynthesis KEGG pathway, map 00941 (https://www.genome.jp/kegg-bin/show_pathway?map00941) (S6 Fig). According to our transcriptome profile results, two genes, *Os06g0626700* (*OsINS*) and *Os01g0372500* (*OsANS*), were upregulated and one gene *Os04g0630800* (*OsANR*) was down regulated in NIL compared with 9311 (S6 Fig, blue rectangle). *OsANS* was also upregulated in OE compared with Nipponbare (Fig 6). Therefore, we tested *OsINS* and *OsANR* in OE lines and Nipponbare. Consistent with expression profile in NIL and 9311, the *OsINS* was significantly upregulated and the *OsANR* was downregulated in OE lines compared with Nipponbare, indicated that *OrC1* regulated some of the same genes in *indica* and *japonica*background.

### Haplotype analysis

The promoter and coding regions of *C1* alleles from 180 rice accessions were isolated and analyzed. The haplotype analysis of promoter region showed that the variation in this region was not associate with coloration traits (S7 Fig). For coding region, 12 haplotypes (H_1 to H_12) were identified. Among them, H_2, 9, 10, 12 were functional, and H_1, 3, 4-8, 11 were non-functional with different deletions (Fig 8a). Only two *O*. *rufipogon* and seven *O*. *nivara* have non-functional allele. The combination of the phenotypes and genotypes of the association panel showed that 24 cultivated rice accessions with non-functional allele showed at least one coloration trait (purple hull or stigma), indicating the presence of other determinant factors besides *C1*. All *indica* and wild rice in H_2 displayed purple apiculus, leaf sheath and stigma, almost all *japonica* in H_2 only displayed purple apiculus (S7 Table). Haplotype network analysis showed that the most frequent non-functional alleles in *indica* and *japonica* are different, which were H_1 and H_7 in *indica* and *japonica*, respectively (Fig 8b), indicating multiple origins of *C1* in two subspecies.

**Fig 8.**
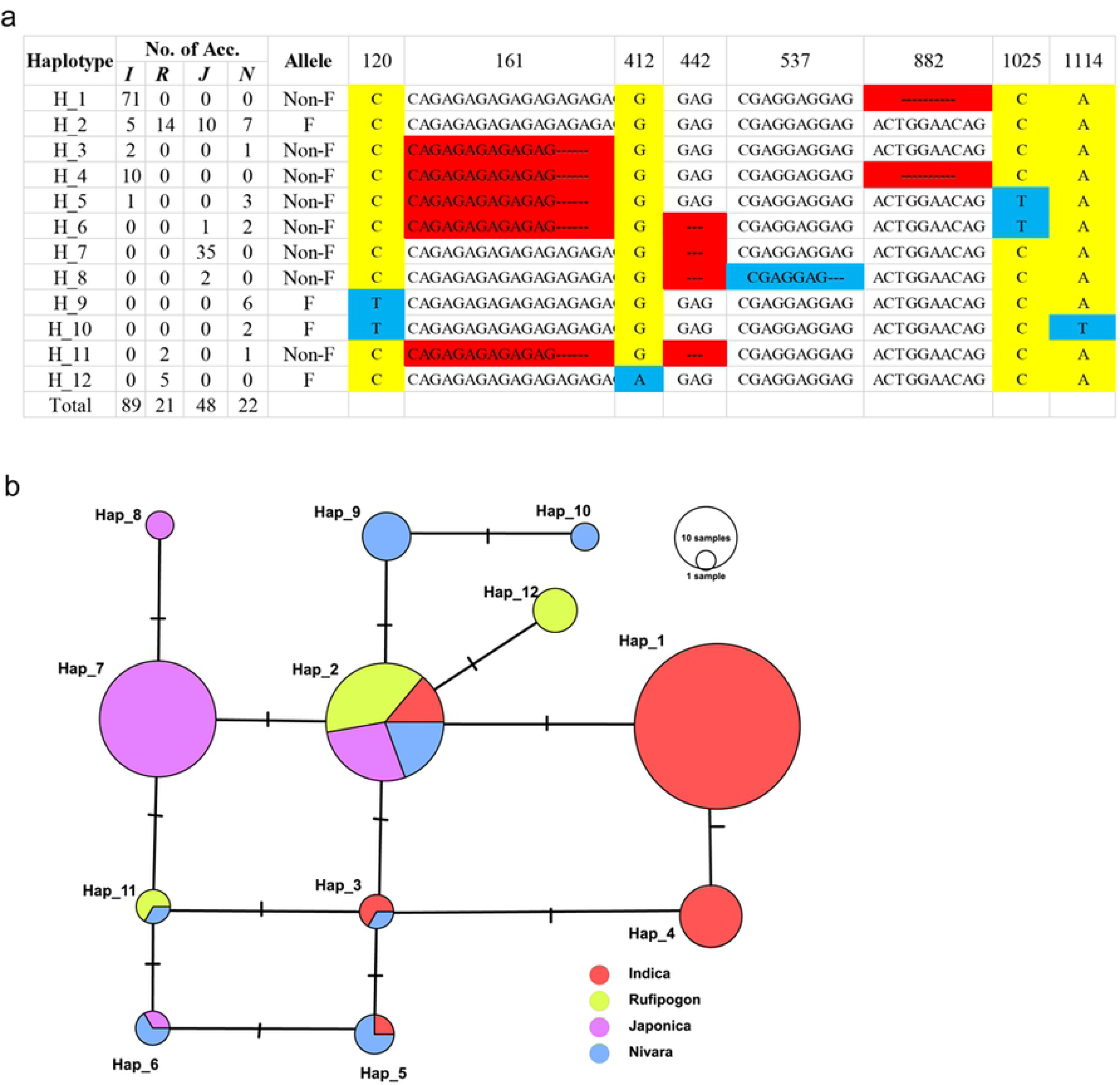
Genotype analysis of coding region of *C1*. (a) Sequence polymorphism of different haplotypes of *C1* in 180 accessions. No. of Acc.: number of accessions, I: *indica*, J: *japonica*, R: *O. rufipongon*, N: *O. nivara*, F: functional, Non-F: non-functional. (b) Haplotype network of *C1*.

## Discussion

Although the spatiotemporal regulation of anthocyanin biosynthesis has been well documented and the chromogen *C* gene, a major coloration gene, has also been fine mapped in cultivated rice [20, 32, 33], the allele in wild rice or full functional allele of *C* gene in rice has not been cloned. Our study identified a functional *C* gene, which encodes a R2R3-MYB transcription factor in wild rice. Given the high heterogeneity in the genome of *O. rufipogon*, it is difficult to clone novel genes which have been lost or weakened in cultivated rice during domestication. CSSLs from inter specific hybridization represent a powerful and useful genetic resource for genome research [34]. Fine mapping was employed firstly in search of anthocyanin genes using a set of CSSLs [29]. The CSSLs in this study has been testified to be an excellent platform for large-scale gene discovery in wild rice [35–37].

Anthocyanin biosynthesis is affected by a variety of environmental factors, including light, temperature, and fertilization [38, 39]. In order to eliminate the environmental effects on coloration, the purple coloration of CSSL population was detected under five environments. A stablely expressed QTL associated with purple coloration was fine mapped on same location of cultivated rice *OsC1*, which was located on chromosome 6 by GWAS [28], hence we named this gene *OrC1*. According to the phylogenetic analysis and sequence blast with homologs in *indca* 9311 and *japonica* Nipponbare, *OrC1* is a complete allele of *C* gene, encoding a R2R3 MYB transcriptor. Although the other organs of NIL such as stem and leaf have no purple coloration, the *C1* gene constitutively expressed in all tissues, and has consistent expression patterns in both NIL and 9311 (Fig 3b). We also sequenced the promoter region of *C1* gene in 180 rice accessions, no haplotype was associated with purple traits (S7 Fig), indicating that the anthocyanin biosynthesis in rice was mostly controlled by C1 protein function but not at gene expression level. As domestication related gene, we deduced that the artificial selection occurs in its coding region but not the promoter region.

Anthocyanin biosynthesis in different rice tissues is primarily controlled by R2R3-MYB transcription factors and the ternary MBW complexes, which comprised of R2R3-MYBs, bHLHs and WD40 [40, 41]. In this study, the NIL-*OrC1* in *indica* background shown a co-segregated purple leaf sheath, stigmas and apiculus, the purple coloration is completely related and shown a 3:1 segregation in F2 population of NIL/9311, which indicated that the coloration of three tissues was controlled by a single dominant gene *OrC1*. However, the OE lines of *OrC1* in *japonica* background only had a purple apiculus but non-colored leaf sheath and stigmas. The different purple traits in NIL and OE are due to their different MBW complex. At least six bHLH genes and three WDR genes have non-synonymous SNP between 9311 and Nipponbare (S5 Fig). The MBWs containing these bHLH or WD40 might have different functions in anthocyanin biosynthesis in different tissues. Moreover, both 9311 and Nipponbare have a non-functional allele of *Os04g0557500* (S5 Fig), which was reported as a tissue regulator and controls pigmentation in hull [11]. Hence, both NIL and OE accumulated pigments only in apiculus but not the hull. Previously studies demonstrated that *OsC1* is the determinant factor of anthocyanin biosynthesis in leaf sheath and apiculus [25, 26], but has not been reported to be involved in pigmentation in stigmas. It is likely that an MBW complex consisting of *OrC1* and a tissue-specific regulator regulated the anthocyanin biosynthesis in stigmas of NIL. Further work is needed to determine the exact bHLH TF combined with *OrC1*. Moreover, haplotype analysis showed that the functional *C1* allele led to anthocyanin accumulation in three tissues in *indica* and only in apiculus in *japonica* (Fig 8 and S7 Table), consistent with the results of *OrC1* function in 9311 and Nipponbare (Fig 2 and Fig 6). Besides *C1*, other redundant genes must be involved in the regulation of anthocyanin biosynthesis in rice stigma.

Transcriptome profiles revealed that almost all the structural genes in the flavonoid biosynthesis pathway were induced by *OrC1* in NIL, but only *ANS*, *CHI*, and *F3H* were induced in OE (Fig 4 and Fig 6). *OrC1* or MBW complex regulate different branches of flavonoid biosynthesis pathway by activating a subset of structural genes. In rice, structure genes are activated by MYB and MBW with redundancy [41], we believe that some bHLH TFs of 9311 is functional and some of Nipponbare is non-functional, the coloration of NIL was due to both OrC1 and MBW complex, but the purple apiculus of OE lines was only caused by OrC1 protein. Furthermore, *OrC1* is involved not only in flavonoid and phenylpropanoid (a basic skeleton of flavonoids) biosynthesis, but also in singling pathways like MAPK, and regulated bHLH and WD40 genes (Fig 5). This indicates that *C1* gene is involved in other physiological processes besides anthocyanin biosynthesis.

The exact metabolites of anthocyanin that *C1* gene produced is still not clear. Shin et al. (2006) transformed maize *C1* and *R-S* regulatory genes into rice using endosperm specific promoters, the produced flavonoids mostly are anthocyanins including dihydroquercetin (taxifolin), dihydroisorhamnetin (3′-O-methyl taxifolin) and 3′-O-methyl quercetin [42]. We have detected six DAMs accumulated in both NIL and OE line (S6 Table). Because we only tested the anthocyanin metabolites, so other flavonoids metabolites are not excluded. Three genes involved in the common DAM biosynthesis pathway have the same expression pattern in NIL_vs_9311 and OE_vs_Nipponbare. Take together the seven structural gene expression profiles, we deduced that *OrC1* regulated *OsCHI*, *OsF3H*, *OsANS*, *OsINS*, and *OsANR* independently from the MBW complexes. These results set a foundation to understand the regulatory mechanisms of *OsC1* in the anthocyanin biosynthesis pathway.

The *C1* involved anthocyanin biosynthesis pathway pre-exists in wild rice but is absent in most cultivated rice, indicating a strong negative human selection of this trait. Haplotype network analysis showed that the functional mutations of *C1* had multiple origins and been selected independently in two subspecies (Fig 8). There are two main hypotheses for the cultivated rice domestication: single origin and multiple origin [43]. Some well-documented domestication genes of rice, such as *sh4* for seed shattering [44], and *prog1* for erect growth [45], supporting the single-origin hypothesis. Our haplotype analysis of *C1* supports multiple origin. More interestingly, the non-functional *C1* allele is much closer to *O. nivara* than *O. rufipongon*, which supports the theory that the origin of cultivated rice is *O. nivara*, an annual wild rice species that is generally considered an intermediate between *O. rufipongon* and cultivated rice; however, the direct ancestor of cultivated rice remains controversial.

## Materials and Methods

### Plant materials

A set of CSSLs produced from common wild rice (*O. rufipogon*) as the donor and an elite *indica* variety, 9311, as the recurrent parent was developed in our laboratory [29]. The CSSLs and 9311 were grown under five environmental conditions as shown in S1 Table. A panel of 180 rice accessions (S7 Table) including 89 *O*. *sativa indica*, 48 *O*. *sativa japonica*, 21 *O*. *rufipogon* and 22 *O*. *nivara*, was selected from a natural rice germplasm population that is maintained in our laboratory. The purple traits were recorded at two environments at Beijing and Sanya, respectively.

### Sequence and phylogenetic analyses

The predicted amino acid sequences of C1 in *O. rufipongon* and MYB homologous proteins in other species were downloaded from NCBI (http://phytozome.jgi.doe.gov/pz/portal.html). Multiple sequence alignments were performed with DNAMAN software. A phylogenic tree was constructed using the Maximum Likelihood method and software MEGA version 5.1 [46]. The de novo genomic sequencing of the panel of 180 rice accessions was performed by our laboratory previously. *C1* gene genomic sequence were isolated from the genomic sequencing data. The SSR primers used in this study were previously published [47], InDel primers were designed in our laboratory. The other primers used in this study were designed online at the NCBI website (https://www.ncbi.nlm.nih.gov/) based on the 9311 reference genome sequence. Alignments were performed on the Gramene (http://www.gramene.org/) website to ensure the accuracy of the location and the specificity of the primers. Sequences of all primers used in this study are shown in S2 Table.

### Subcellular localization

The full-length coding region of *OrC1* was inserted into the PAN580-GFP plasmid to generate 35S::OrC1-GFP constructs and transformed into rice protoplasts and tobacco leaves. The GFP fluorescence in rice protoplasts and leaf epidermal cells were observed using a laser scanning confocal microscope (LSM880, Leica)

### RNA extraction and real time PCR

Total RNA was extracted from different rice tissues using the RNA RNeasy Plant Mini Kit (Qiagen, Beijing, China), followed by treatment with RNase-free DNase (TaKaRa, Dalian, China) to remove genomic DNA contamination. qRT-PCR was performed using an ABI 7500 real time PCR system (Applied Biosystems) following the manufacturer’s instructions. The poplar endogenous control gene (Actin) was used as an internal control. Gene-specific primers are listed in S2 Table. Relative expression was calculated using the 2^-[Δ][Δ]Ct^ method [48]. Each sample was amplified in triplicate.

### Transcriptome analysis

The leaf sheath of 15-day seedlings of NIL-*OrC1* and 9311 were collected for RNA extraction. RNA samples were sent to Genedenovo Biotechnology Co., Ltd (Guangzhou, China) for RNA sequencing. The rRNA removed reads of each sample were then mapped to reference genome by TopHat2 [49]. The reconstruction of transcripts was carried out with Cufflinks (http://cole-trapnell-lab.github.io/cufflinks/install/) and TopHat2. Principal component analysis was performed with R package Rmodest (http://www.r-project.org/). Data analyses, including DEGs, GO and pathway enrichment, were performed by Genedenovo Biotechnology Co., Ltd.

### Rice transformation

*Agrobacterium*-mediated transformation was performed by Biorun Biosciences Co., Ltd (Wuhan, China) following their standard procedures and protocols.

### Anthocyanin metabolite determination

The leaf sheath of 15-day seedlings of NIL, 9311, OE lines and Nipponbare, the upper one third of hull at 10 days after flowering of OE and Nipponbare were collected. The sample extracts were analyzed using an LC-ESI-MS/MS system (HPLC, Shim-pack UFLC SHIMADZU CBM30A system, http://www.shimadzu.com.cn/; MS, Applied Biosystems 4500 Q TRAP, http://www.appliedbiosystems.com.cn/). The sample preparation, extract analysis, metabolite identification and quantification were performed at Wuhan MetWare Biotechnology Co., Ltd. (http://www.metware.cn) following their standard procedures and were previously described in details by [50–52]

## Acknowledgements

We especially thank Dr. Peng Zhang (The University of Sydney, Australia) and Dr. Hao Chen (Huazhong Agricultural University, China) for critical reading of the manuscript. This work was supported by the National Key R&D Program of China (2016YFD0100101) and the National Natural Science Foundation of China (31471471).

## Author contributions

WH Qiao, JH Lan and QW Yang conceived and designed the experiments. WH Qiao and YY Wang performed the experiments and wrote the paper. R Xu, ZY Yang, JR Wang and JF Huang analyzed the data. Y Sun, L Su and LZ Zhang performed fine mapping. SJ Liu, YL Tian, LM Chen and X Liu contributed to field investigation. All authors read and approved the final manuscript.

## Supporting information

**S1 Table. The rice crop locations used in this experiment**

**S2 Table. Informations of the primers used in this study**

**S3 Table. The segregation of purple coloration traits in the CSSL52/9311 F_2_ population**

**S4 Table. Predicted open reading frames in the mapped location**

**S5 Table. DEGs in the trancriptome profile of 9311_vs_NIL**

**S6 Table. Differentially accumulated anthocyanins between 9311_vs_NIP and Nipponbare_vs_OE**

**S7 Table. Phenotypes of coloration of a panel of rice accessions**

**S1 Fig. Principal component analysis of the trancriptome samples.** P93-11 and OrC1 represent those for 9311 and NIL, respectively, One replicate of NIL that is clearly different from other two was deleted.

**S2 Fig. Gene ontology enrichment of DEGs in leaf sheath of NIL and 9311at the seedling stage.**

**S3 Fig. Regression analysis between the results of RNA-Seq and qRT-PCR experiments.**

**S4 Fig. Identification of overexpression lines of *OrC1*.** 35S:OrC1 was transformed into Nipponbare by an *Agrobacterium*-mediated transformation. (a) Ten transgenic lines was confirmed by PCR analysis using genomic DNA with gene-specific primers. (b) Ten positive transgenic lines were analyzed by qRT-PCR to determine the OrC1 expression level. Nip:Nipponbare, OE: overexpression line.

**S5 Fig. Comparison of bHLH and WDR genes between Nipponbare and 9311.** The genomic sequence of each gene was from http://oge-ensembl.gramene.org/index.html, red rectangles indicate non-synonymous SNPs.

**S6 Fig. KEGG pathway map of favonoid biosynthesis in 9311_vs_NIL transcriptome profile.** The red and green rectangles indicate up and down regulation, respectively. Anthocyanin biosynthesis pathway was marked using a blue rectangle, ID of the three genes in each pathway were showed inside the blue rectangle, the number in each bracket behind ID represents log2(FC).

**S7 Fig. Genotype analysis of promoter region of *C1*.** (a) Sequence polymorphism of different haplotypes of *C1* promoter from 180 accessions. No. of Acc.: number of accessions, I: *O*. *sativa. indica*, J: *O*. *sativa. japonica*, R: *O*. *rufipongon*, N: *O*. *nivara*. (b) Haplotype network of *C1* promoter.

